# Representational models: A common framework for understanding encoding, pattern-component, and representational-similarity analysis

**DOI:** 10.1101/071472

**Authors:** Jörn Diedrichsen, Nikolaus Kriegeskorte

**Affiliations:** Brain and Mind Institute, Department for Computer Science, Department for Statistical and Actuarial Science, Western University, Canada; Cognitive and Brain Sciences Unit, Cambridge University, UK

## Abstract

Representational models specify how activity patterns in populations of neurons (or, more generally, in multivariate brain-activity measurements) relate to sensory stimuli, motor responses, or cognitive processes. In an experimental context, representational models can be defined as hypotheses about the distribution of activity profiles across experimental conditions. Currently, three different methods are being used to test such hypotheses: encoding analysis, pattern component modeling (PCM), and representational similarity analysis (RSA). Here we develop a common mathematical framework for understanding the relationship of these three methods, which share one core commonality: all three evaluate the second moment of the distribution of activity profiles, which determines the representational geometry, and thus how well any feature can be decoded from population activity with any readout mechanism capable of a linear transform. Using simulated data for three different experimental designs, we compare the power of the methods to adjudicate between competing representational models. PCM implements a likelihood-ratio test and therefore provides the most powerful test if its assumptions hold. However, the other two approaches – when conducted appropriately – can perform similarly. In encoding analysis, the linear model needs to be appropriately regularized, which effectively imposes a prior on the activity profiles. With such a prior, an encoding model specifies a well-defined distribution of activity profiles. In RSA, the unequal variances and statistical dependencies of the dissimilarity estimates need to be taken into account to reach near-optimal power in inference. The three methods render different aspects of the information explicit (e.g. single-response tuning in encoding analysis and population-response representational dissimilarity in RSA) and have specific advantages in terms of computational demands, ease of use, and extensibility. The three methods are properly construed as complementary components of a single data-analytical toolkit for understanding neural representations on the basis of multivariate brain-activity data.

**Author Summary**

Modern neuroscience can measure activity of many neurons or the local blood oxygenation of many brain locations simultaneously. As the number of simultaneous measurements grows, we can better investigate how the brain represents and transforms information, to enable perception, cognition, and behavior. Recent studies go beyond showing *that* a brain region is involved in some function. They use representational models that specify *how* different perceptions, cognitions, and actions are encoded in brain-activity patterns. In this paper, we provide a general mathematical framework for such representational models, which clarifies the relationships between three different methods that are currently used in the neuroscience community. All three methods evaluate the same core feature of the data, but each has distinct advantages and disadvantages. Pattern component modelling (PCM) implements the most powerful test between models, and is analytically tractable and expandable. Representational similarity analysis (RSA) provides a highly useful summary statistic (the dissimilarity) and enables model comparison with weaker distributional assumptions. Finally, encoding models characterize individual responses and enable the study of their layout across cortex. We argue that these methods should be considered components of a larger toolkit for testing hypotheses about the way the brain represents information.

## Introduction

The measurement of brain activity is rapidly advancing in terms of spatial and temporal resolution, and in terms of the number of responses that can be measured simultaneously [1]. Modern electrode arrays and calcium imaging enable the recording of hundreds of neurons in parallel. Electrophysiological signals that reflect summaries of the population activity can be recorded using both invasive (e.g. the local field potential, LFP) and non-invasive techniques (e.g. scalp electrophysiological measurements) at increasingly high spatial resolution. Modern functional magnetic resonance imaging (fMRI) enables us to measure hemodynamic activity in hundreds of thousands of voxels across the entire human brain at sub-millimeter resolution.

In order to translate advances in brain-activity measurement into advances in computational theory [2], researchers increasingly seek to test representational models that capture both what information is represented in a population of neurons, and how it is represented. Knowing the content and format of representations provides strong constraints for computational models of brain information processing. We refer to hypotheses about the content and format of brain representations as *representational models*, and address here the important methodological question of how to best test such models.

Referring to an activity pattern as a “representation” constitutes a functional interpretation [3], which requires not only that the represented variable (such as a perceptual property, some cognitive content, or an action parameter) is encoded in the pattern of activity in a format that can be read out by downstream neurons, but also that the information is actually used by other brain regions and, thus, *serves a functional purpose* [4]. The representational interpretation therefore ultimately needs to be supported by evidence for a cause-and-effect relationship between the activity and downstream neural and behavioral responses. Testing causal effects of activity patterns is beyond the scope of the present paper. However, we note that a good brain-computational model must, as a necessary condition, be able to explain the information present in brain regions involved in task performance and the format in which this information it is encoded.

For a population code to constitute an *explicit representation*, another area must be able to read out the represented variable directly using a neurobiologically plausible readout mechanism, such as linear or radial-basis-function decoding [2, 5, 6]. Note that this definition of explicit does not restrict us to highly localized codes, such as the “grandmother neuron” [7], but encompasses widely distributed codes.

An example of an implicit representation is the representation of object category in the retina. The retina clearly contains information about object category, and an aspect of its function is to convey this information. However, it does not *explicitly represent* object category. Multiple stages of nonlinear tranformation along the ventral visual stream are required to render the category of an object explicit. Inferior temporal cortex contains a representation of object category [8, 9], along with representations of much additional information [10].

Many researchers have used linear decoding methods to reveal explicit information in neural representations [11–13]. Representational models, as considered here, go one step further: they fully characterize the representational geometry, defining all explicitly represented features in a region, how strongly each of them is represented (signal to noise ratio), and how the activity patterns associated with different features relate to each other. Representational models therefore fully specify the explicit representational content.

To define representational models formally, we need to consider two complementary perspectives on activity data, as illustrated in Fig. 1. The activity of many neurons, or more generally *measurement channels* (neurons, electrodes, or fMRI voxels), can be measured across a range of *experimental conditions* (stimuli, movements, or tasks). Thus, each channel will have an *activity profile,* which can be plotted as a point in the space spanned by the experimental conditions (Fig. 1b). A representational model specifies a probability distribution of activity profiles in the space spanned by the experimental conditions. It treats the true activity profiles as a random variable and predicts, for each possible activity profile, the probability of observing a measurement channel exhibiting that profile. It does not predict the activity profile for each individual channel actually measured. The motivation for this approach derives from the idea that the computational function of a region does not depend on specific neurons having specific response properties, but on the fact that certain features can be read out from the population by downstream neurons. The probability distribution over activity profiles determines which features can be linearly read out from the code and the signal-to-noise ratio of the readout. By basing further analyses on the probability distribution of the activity profiles, we are disregarding three aspects of the code: (1) which neuron fulfills which function, (2) where neurons are located within a cortical area, and (3) the degree to which the information about a given represented feature is concentrated in a few neurons (as in single-cell selectivity for a represented feature) or spread out over the population. Ignoring these aspects may be viewed as an advantage or a disadvantage, depending on the level of description that a researcher is interested in. We argue that treating activity profiles as random vectors is a simplification that is useful for drawing computational insights from population activity measurements.

**Fig 1.**
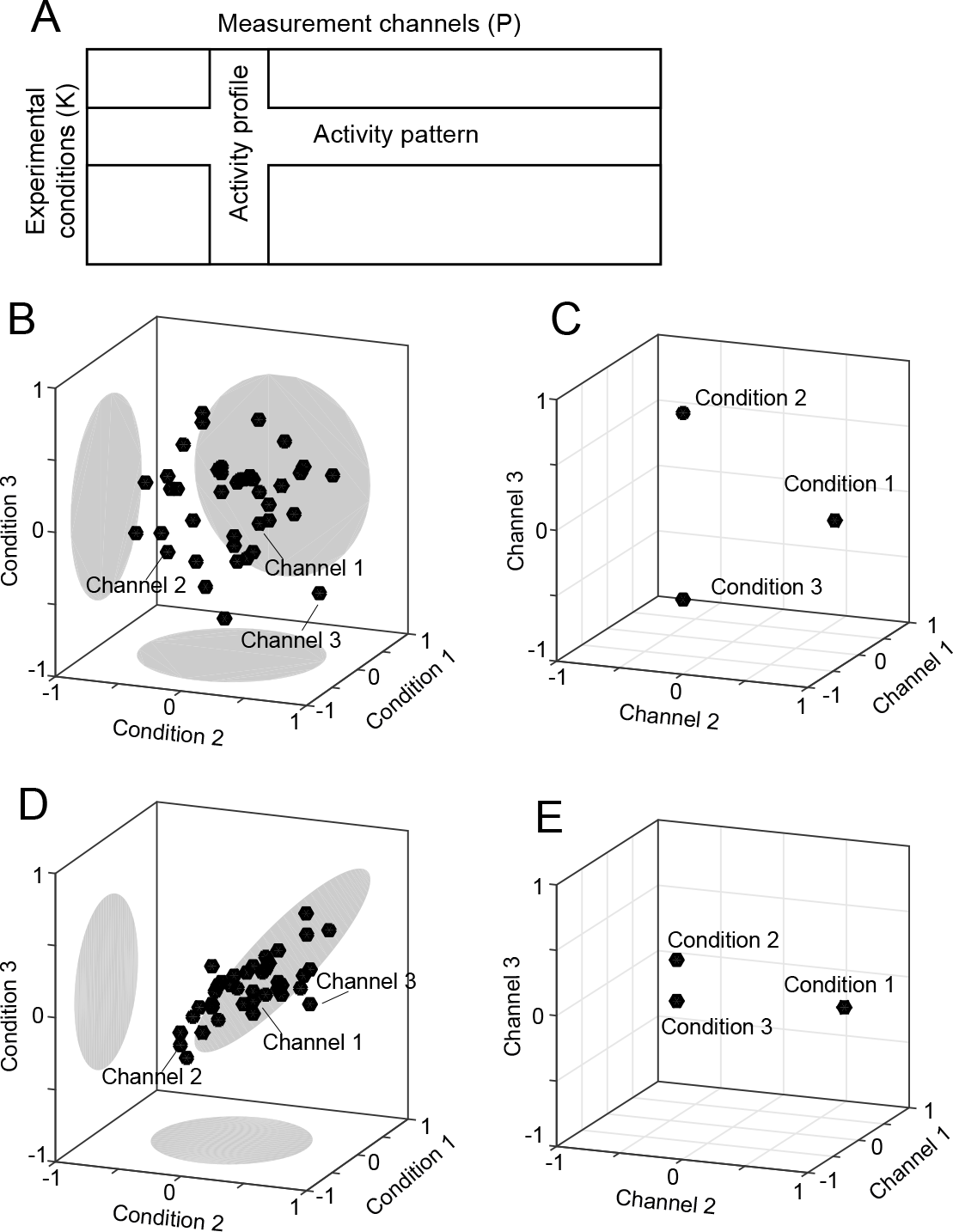
Two complementary perspectives on population activity. (**A**) The multivariate activity data can be viewed as a set of activity profiles (columns) or as a set of activity patterns (rows). An activity profile is a vector of responses of a single channel across experimental conditions. An activity pattern is a vector of responses across all channels for a single condition. Activity data can be visualized by plotting activity profiles as points in a space defined by the experimental conditions (B,D), or by plotting the activity patterns as points in a space defined by the measurement channels (C,E). (**B**) If the activities are uncorrelated between conditions, then (**C**) the corresponding activity patterns of all three conditions are equidistant to each other, and can be equally well distinguished. (**D**) If the activities are positively correlated for two conditions that elicit similar regional-mean activation (conditions 2 and 3 here), then (**E**) the activity patterns for these conditions are closer to each other and can be less well distinguished.

In this paper, we show that the multivariate *second moment* of the activity profiles fully defines the representational geometry and with it all the information that can linearly or nonlinearly decoded. In particular, the second moment determines the signal-to-noise ratio with which any feature can be linearly decoded. We discuss three established methods for adjudicating between representational models: encoding analysis, pattern-component modeling (PCM) and representational similarity analysis (RSA, see Table 1). We show that these three techniques all exclusively rely on information contained in the second moment. This core commonality enables us to consider these methods in the same formal framework.

**Table 1.** Comparison of encoding analysis with regularization, pattern component modelling (PCM), and representational dissimilarity analysis (RSA).

|  | Encoding analysis | PCM | RSA |
| --- | --- | --- | --- |
| Model definition | Model-feature matrix $\mathbf{M}$, regularization / prior | Predicted second-moments matrix ($\mathbf{G}$) | Representational dissimilarity matrix (RDM) |
| First-level parameters (characterizing individual activity profiles) | One weight per feature and measurement channel | None; integrated out in the likelihood | None; integrated out when calculating dissimilarities |
| Second-level parameters (characterizing the distribution of activity profiles) | Regularization / Ridge coefficient (determined by noise / signal ratio) | Scale parameter $s$, Noise variance | Scaling between predicted and observed distances ($s$) |
| Prediction target | Responses to test conditions | Distribution of measurement channels in activity-profile space | Dissimilarities among activity patterns |
| Training data required | always | not for fixed models, only if additional second-level parameters are to be fitted | not for fixed models, only if additional second-level parameters are to be fitted |
| Explicit likelihood for fitting additional model parameters | No – need to do nested within crossvalidation | Yes | Yes |
| Fitting algorithms for model parameters | - | EM<br>Gradient descent<br>Newton-Raphson | Linear and non-negative regression<br>IRLS |

In *encoding analysis* [14, 15], representational models are defined in terms of the underlying *features* (Fig. 2A). Each activity profile can be characterized by a linear combination of such features. Examples include Gabor filters [16] (for a low-level visual representation), abstract semantic dimensions [17] (for a cognitive representation), and force, direction or hand position [18–20] (for a movement representation). The importance of each feature in each channel is measured by a feature weight. Feature weights are considered first-level parameters in our framework, as they describe the individual activity profiles, as opposed to second-level parameters that describe the distribution of the activity profiles (Table 1). The large number of parameters (number of features in the model times number of channels in the measurements) engenders a danger of overfitting. Encoding models are therefore commonly evaluated using cross-validation: The feature weights are estimated on a training set, and the model is evaluated in terms of its performance at predicting left-out data [14]. The test data may consist in a sample of experimental conditions not used in training, so as to test the model’s generalization performance [15, 16]. While many studies use simple linear regression to estimate the weights [15, 21], it is increasingly common to use a regularization penalty (for example the L2 norm of the vector of weights) [16, 17]. We will show that regularization is not merely a technical trick used in fitting a given model. Instead, the regularization (and its implicit distributional assumptions) are an essential part of the representational hypothesis that is tested. Without it, encoding models do not specify a probability distribution with a finite second moment and thus do not define the linear decodability of different features.

**Fig 2.**
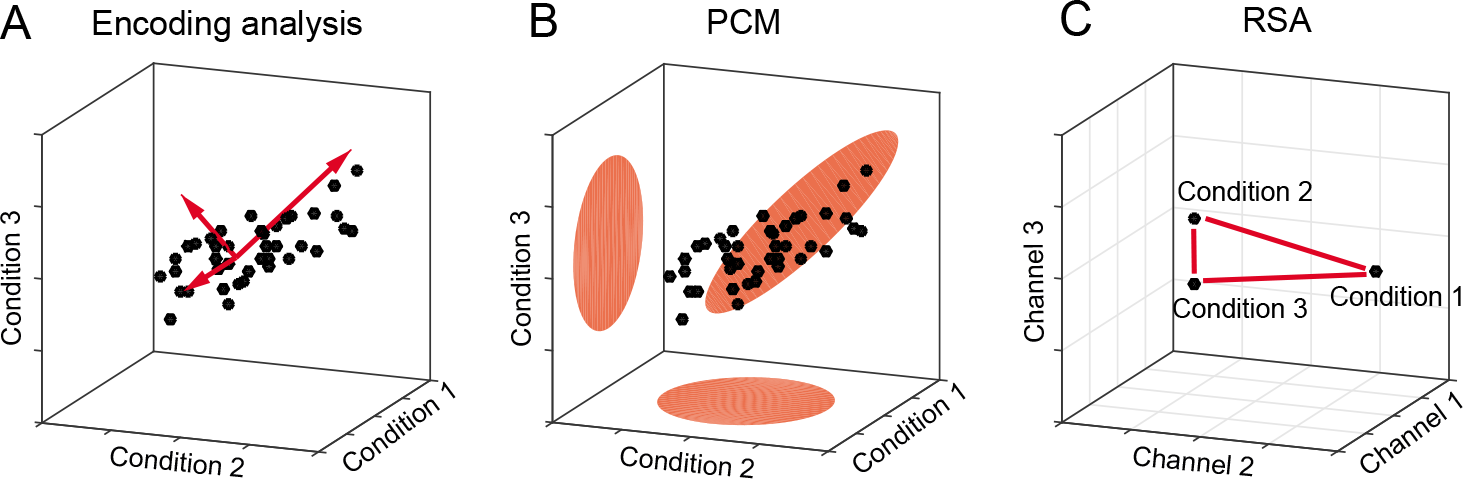
Three approaches to testing representational models. (A) In encoding analysis, the distribution of activity profiles is described by the underlying features (red vectors). The direction of a feature vector determines the associated activity profile, and the length the strength of the feature encoding in the representation. (**B**) PCM models the distribution of the activity profiles as a multivariate Gaussian. This model is parametrized by the second moment of the activity profiles, which determines at what signal-to-noise ratio any feature is linearly decodable from the population. (**C**) RSA uses the representational distances (or, more generally, dissimilarities) between activity patterns as a summary statistic to describe decodability and hence the second moment of the underlying distribution.

*Pattern component modeling* [22] is based on an explicit generative model of the process that produced the data and can be considered a Bayesian approach. The true activity profiles are assumed to have a multivariate Gaussian distribution in the space spanned by the experimental conditions (Fig. 2B). This formulation enables us to evaluate the marginal likelihood of the observed activity profiles under the probability distribution specified by the model. Thus, we do not fit any first-level parameters (feature weights) and hence reduce the risk of overfitting. This enables us to compare models with different numbers of features without having to correct for model complexity. If the assumptions of the generative model hold, PCM implements the likelihood-ratio test between models [23], which by the Neyman-Pearson lemma [24], is the most powerful test of its size. In theory, therefore, PCM should yield more accurate inferences than any of its competitors, that is it should be able to more sensitively adjudicate among competing models.

Finally, *representational similarity analysis* (RSA [9, 25, 26]) approaches the problem from a complementary perspective. Rather than considering the activity profiles of the measurement channels as points in the space spanned by the conditions (Fig. 1B,D), it considers the activity patterns associated with the experimental conditions as points in the space spanned by the measurement channels (Fig. 1C,E). RSA then uses the representational distances (Fig. 2C) between the conditions as a summary statistic. We will see that these distances again exclusively depend on the second moment of the distribution of activity profiles. Having obtained a matrix of dissimilarities between activity patterns (the representational dissimilarity matrix, RDM), RSA then tests models by comparing the observed distances to the distances predicted by each representational model. This can be done by calculating rank-based correlations [27] or Pearson correlations [28]. Here we show that for near-optimal inferences it is important to take the co-dependence structure of the distance estimates into account, for example by using a multivariate normal approximation to the joint distribution of the cross-validated Mahalanobis distances [29, 30].

In the remainder of the paper, we first introduce the second moment of the activity profiles and explain why it is the sufficient statistic of the representational geometry and thus of linear and nonlinear decodability. We then define the three methods in detail, and show how they related to the second moment. Finally, using simulated data and models taken from our fMRI work, we assess the statistical efficiency, i.e. how well these methods adjudicate between two or more competing representational models given limited data. We also compare the methods in terms of their computational efficiency.

## Materials and Methods

### Basic definitions

All symbols used in the following derivations are summarized in Table 2. First, we define **U** to be the matrix of noiseless activity profiles with *K* (number of experimental conditions) rows and *P* (number of measurement channels) columns. Each row of this matrix is an activity pattern, the response of the whole population to a single condition. Each column of this matrix is an activity profile (Fig. 1A).

**Table 2.** Notation used. For non-scalars, the second column indicates the vector / matrix size.

|  |  |  |
| --- | --- | --- |
| $K$ | | Number of conditions |
| $M$ | | Number of independent partitions (imaging runs) |
| $P$ | | Number of measurement channels (voxels, electrodes, neurons) |
| $N$ | | Overall number of measurements ($N_m \times M$) |
| $Q$ | | Number of features in model |
| $\mathbf{U}$ | $K \times P$ | Matrix of true activation patterns |
| $\mathbf{u}_{i,\cdot}$ | $1 \times P$ | Activation pattern for condition $i$; $i^{\text{th}}$ row of $\mathbf{U}$ |
| $\mathbf{u}_{\cdot,j}$ | $K \times 1$ | Activation profile for measurement channel $j$; $j^{\text{th}}$ column of $\mathbf{U}$ |
| $\hat{\mathbf{U}}^{(m)}$ | $K \times P$ | Matrix of estimated activity patterns, based on data from partition $m$ |
| $\tilde{\mathbf{U}}^{(\sim m)}$ | $K \times P$ | Model prediction for activity patterns, based on data independent of $m$ |
| $\mathbf{M}$ | $K \times Q$ | Matrix of model features for all condition |
| $\mathbf{W}$ | $Q \times P$ | Matrix of voxel weights for each feature |
| $\mathbf{Y}$ | $N \times P$ | Matrix of brain measurements, concatenated activity estimates or time series data |
| $\mathbf{Z}$ | $N \times K$ | Design matrix, indicating how measurements relate to activity patterns |
| $\mathbf{X}$ | $N \times R$ | Design matrix containing $n$ regressors of no-interest |
| $\mathbf{G}$ | $K \times K$ | Second moment of $\mathbf{U}$ |
| $d_{i,k}$ | | Distance between condition $i$ and $k$ |
| $J$ | | Number of distances, normally $K(K-1)/2$ |
| $\mathbf{D}$ | $K \times K$ | Representational dissimilarity matrix of all pairwise distances |
| $\mathbf{d}$ | $J \times 1$ | Vector of all pairwise distances |
| $\tilde{\mathbf{d}}$ | $J \times 1$ | Vector of predicted distances |
| $\mathbf{C}$ | $J \times K$ | Contrast matrix, defining the $J$ pairwise differences between conditions |
| $\Sigma_P$ | $P \times P$ | Variance-covariance matrix between the $P$ voxels |
| $\Sigma_K$ | $K \times K$ | Variance-covariance matrix of the columns of $\hat{\mathbf{U}}^{(m)}$ |
| $\mathbf{V}$ | $N \times N$ | Variance-covariance matrix of $\mathbf{Y}$ |
| $\mathbf{S}$ | $J \times J$ | Variance-covariance matrix of all pair-wise distances |

Because we are interested in the distribution of activity profiles, but not in the activity profiles per se, we consider the columns of **U** to be a random variable. This is an essential step underlying our common framework, which is justified by the fact that, for the purpose of reading out information, the different measurement channels are exchangeable (see introduction). We assume that the activity profiles are repeatedly measured, with the data consisting of *M* independent partitions, each containing at least one activity measurement for each condition and measurement channel. In the context of fMRI, a partition will consist of a separate phase of data acquisition, e.g. a scanner run. The activity estimates Û^(*m*)^ of partition *m* are the true patterns **U** plus noise **E**^(*m*)^. The noise captures both neural trial-by-trial variability of the activity pattern in a single condition, as well as measurement noise.

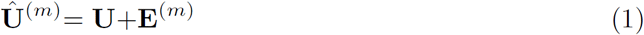

For the purposes of this paper, we assume that the noise is Gaussian, and independent and identically distributed (i.i.d.) across conditions and partitions (homoscedasticity). Possible dependence within each partition, however, can be easily accounted for [29, 31].

### Dependence between measurement channels

The discussion below further assumes that the noise is also i.i.d. across different measurement channels (isotropicity). However, noise in fMRI, MEG, and even invasive electrophysiology exhibits strong correlations between neighboring locations in the brain. To account for these dependencies, we employ multivariate noise normalization (i.e. spatial prewhitening), which has been shown to increase the reliability of inference [32]. Across all measurement channels, we estimate the *P* × *P* variance-covariance matrix across trials, **Σ**_*P*_ and then regularize the estimate by shrinking it towards a diagonal matrix [33]. In the context of fMRI, we can use the residual time series from the fitting of the time-series model to estimate noise covariance [32, 34]. We then post-multiply our activity estimates by 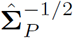, rendering the model errors in the channels approximately uncorrelated. If multivariate noise normalization is not performed or is incomplete, inference will be suboptimal in all three methods (for details see [29]).

### Second moment matrix and linear decodeability

In this section, we show that the *second moment* of the activity profiles fully characterizes the linear decodability of any feature in the space spanned by the experimental conditions. A feature is any property of the experimental conditions, represented as a vector with one entry per condition. The fact that the second moment determines what can be decoded provides a motivation, from the perspective of brain computation, for using the second moment matrix as a summary statistic.

The second moment defines the decodable information, because it determines the representational geometry, i.e. the representational distance matrix. Higher statistical moments may be useful to define the distribution of activity profiles in greater detail (a point we will return to in the Discussion). For example, they capture to what extent particular information is concentrated in single neurons or small sets of neurons – a property that is important to computation if readout neurons cannot integrate information from the entire population that constitutes the code. However, assuming that readout neurons have access to the entire code and can weight activities them in any arbitrary way, the second moment is a sufficient statistic of the decodable information.

The *n^th^* moment of a scalar random variable *u* is *E* (*u^n^*), where *E*() denotes the expected value. Here we use a multivariate extension of the concept, with the second moment of the random vector **u** defined as the matrix *E* (**uu**^*T*^), the expected outer product of the activity profiles, where the expectation is across the measurement channels. The second-moment matrix of the activity profiles is given by

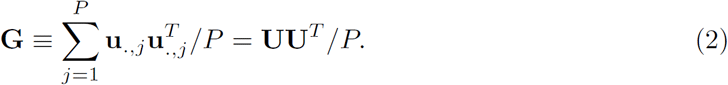

Thus, each cell of this matrix contains the scaled inner product of two activity patterns.

Before calculating **G**, some investigators subtract the mean activity across measurement channels for each condition from the data. In this case, Eq. 2 becomes the variance-covariance matrix of the activity profiles –the second moment around the mean activity profile. Here we do not remove the mean, but use the second moment around zero. From the perspective of a neuron that reads out the activity pattern of an area, any difference between activity patterns across conditions can be used to encode information. Some features (for example, stimulus intensity) may be encoded in the mean activity over all measurement channels. Other properties (for example, stimulus identity) may be encoded in relative activity differences, with some measurement channels responding more to one condition, and others to a different condition. The second moment around zero captures both of these potentially meaningful differences.

Any feature of the conditions that we might want to decode can be defined by a *K* × 1 vector **f** with one entry per condition, which describes how the feature varies across conditions. To obtain a linear read-out estimate 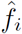 for the feature *f_i_* for a given condition *i*, we weight each channel’s observed activity using the *P* × 1 read-out vector **v**:

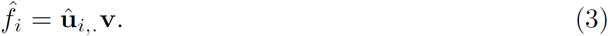

We would like the estimate 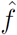 to have very different values for two trials that differ on the feature value, while showing small differences for trials that have the same feature value. We are therefore looking for the readout weight vector **v** that maximizes the ratio between feature variance and error variance, and thus the signal-to-noise ratio (*S*), of the readout:

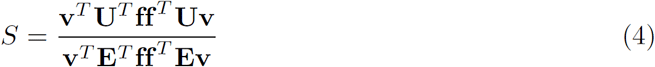

The solution to this equation is commonly known as Fisher’s linear discriminant [35], which, under the assumption of homoscedastic Gaussian noise, is the best achievable linear decoder. If the noise is isotropic (or the data is adequately pre-whitened), then **E**^*T*^ **ff**^*T*^ **E** = **I***b*, where *b* is a constant. The denominator then depends only on the norm of the read-out vector **v**, not on its direction, and can be ignored when **v** is constrained to have a norm of 1. The best readout vector **v** is then given by the first eigenvector of the matrix **U**^*T*^ **ff**^*T*^ **U**, and the quality of the best readout is determined by the corresponding eigenvalue.

Non-zero eigenvalues (*eig*) of a square matrix are invariant to cyclic permutations of the product order:

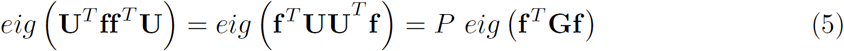

Therefore, the quality of the best linear decoder for *any* feature (as defined by **f**) is fully characterized by the second moment matrix **G** of the pre-whitened activity patterns.

### Representational analysis in the context of fMRI

The methods in this paper were first developed in the context of fMRI data analysis, and our examples will come from this domain. A simple way to apply the analyses to fMRI data is to use as activity estimates (Û^(*m*)^) the regression coefficients, or “beta”-weights, from a first-level time series analysis [36, 37]. The time-series model accounts for the hemodynamic lag and the temporal autocorrelation of the noise. The activity estimates usually express the difference in activity during a condition relative to rest. Activity estimates commonly co-vary together across fMRI imaging runs, because all activity estimates within a partition are measured relative to the same resting baseline. This positive correlation can be reduced by subtracting, within each partition, the mean activity pattern (across conditions) from each activity pattern. This makes the mean of each measurement channel (across condition) zero and thus centers the ensemble of points in activity-pattern space that is centered on the origin.

Rather than using the concatenated activity estimates from different partitions, encoding analysis and PCM can also be applied directly to time series data. As a universal notation that encompasses both situations, we can use a standard linear mixed model [38]:

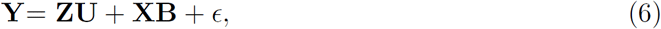
where **Y** is an *N* × *P* matrix of all activity measurements, **Z** the *N* × *K* design matrix, which relates the activity measurements to the *K* experimental conditions, and **X** is a second design matrix for nuisance variables. **U** is the *K* × *P* matrix of activity patterns (the random effects), **B** are the regression coefficients for these nuisance variables (the fixed effects), and **E** is the matrix of measurement errors. If the data **Y** are the concatenated activity estimates, the nuisance variables typically only model the mean pattern for each run. If **Y** consists of time-series data, the nuisance variables typically capture additional effects such as time-series drifts and residual head-motion-related artifacts.

### Representational analysis in the context of neurophysiological recordings

All three methods can also be applied to recordings of single cell activity or neurophysiological potentials [9, 25]. The activity estimates can then be firing rates estimated over a temporal window for each trial, or the power in different frequency bands over time. Because the trial-by-trial variability of firing rates will usually increase with the mean firing rate, it is advisable to use the square root of firing rates to make the data conform better to the assumption that the variance of the noise is independent of the signal [39].

Here we focus on models that treat the activity patterns **U** as static snapshots. To exploit the temporal detail provided by electrophysiological recordings, the analyses can be either performed using a sliding window over the time course of the trial [40–42], or by “stacking” the time series and conditions, resulting in a activity matrix with *TK* rows [43].

### Encoding analysis

An encoding model characterizes the structure of the representation in terms of a set of features [14–17]. We will show in the following that encoding models are representational models as definied by the second moment of the activity profiles. For this to be the case, however, the use of regularized regression is a crticial factor. We will therefore first present the encoding apporach in general, and then show why regularisation is important to test for distributions with a definied second moment.

In general. the value of each feature for each experimental condition is coded in the *model matrix* **M** (*K* conditions by *Q* features). The *feature weight matrix* **W** (*Q* features by *P* channels) then determines how the different model features contribute to the activity profiles of different measurement channels to produce the predicted activity patterns **U**:

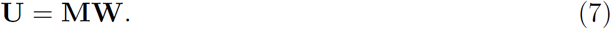

Geometrically, we can think of the features as the basis vectors of the subspace, in which the activity profiles reside (Fig. 2A).

#### Encoding analysis without regularization

To adjudicate among encoding models of different numbers of features – and hence different numbers of free first-level parameters - most researchers use independent test sets [15–17]. A training data set is used to estimate the feature weights for each channel, and the resulting prediction is then evaluated on a held-out test data set. This can be implemented in a statistically efficient manner by using cross-validation, which is usually performed by holding out a single partition (e.g. fMRI imaging run) as a test set, and using the remaining *M*-1 partitions as the training set. Each partition is held out as the test set once and prediction performance is averaged across the *M* folds of cross-validation. Encoding models can also make predictions about conditions that are not in the training set (Discussion). However, we focus our simulations on cases, in which training and test sets include the same experimental conditions.

The weights can be chosen to minimize the sum of squared errors on the training data, i.e. using linear regression:

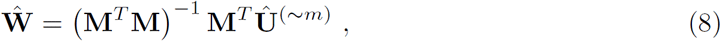
where we define Û^(*∼m*)^ to be the average activity estimates from all partitions except *m*. The prediction for the left-out test data of run *m* is

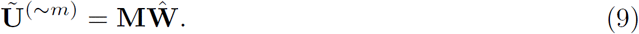

The accuracy of the prediction can be assessed by relating the residual sums-of-squares (SSR) of the prediction to the total sums-of-squares (SST) of the observed activities, summed over all partitions, conditions, and voxels

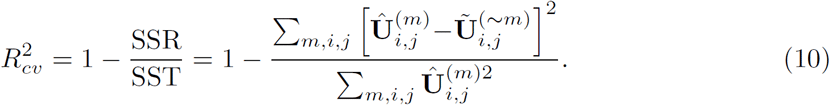

Alternatively, we can evaluate the prediction by correlating the predicted and observed activity patterns across all conditions and channels. Assuming that the mean of each channel across all conditions is zero (given mean pattern subtraction), the correlation is given by

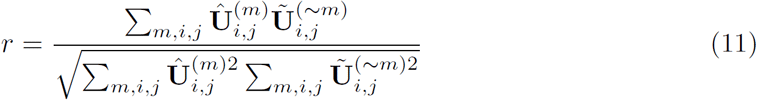

The correlation introduces an arbitrary scaling factor between prediction and observations and, in contrast to Eq. 10, allows the model to over- or under-predict the data by a scalar factor without penalty. Encoding analysis can also be applied directly to the time-series data without an intervening model (Eq. 6).

To understand how encoding analysis adjudicates between models, consider the graphical representation of the estimation process (Fig. 3). In this example, the training data the activity profile of a single measurement channel, which can be visualized as a point in activity-profile space (black cross). Regression analysis can be understood as the orthogonal projection of the measured activity profile onto the linear subspace spanned by the features of the model. The two models depicted in Fig. 3A and Fig. 3B have different features (blue arrows) that define different subspaces (planes with blue outlines). Therefore, the training data is projected onto two different planes and the prediction for the test data differs between the two models. The model with a subspace that better describes the cloud of activation profiles will make better predictions overall across the measurement channels, show lower cross-validation error, and will hence be more likely selected as the winning model.

**Fig 3.**
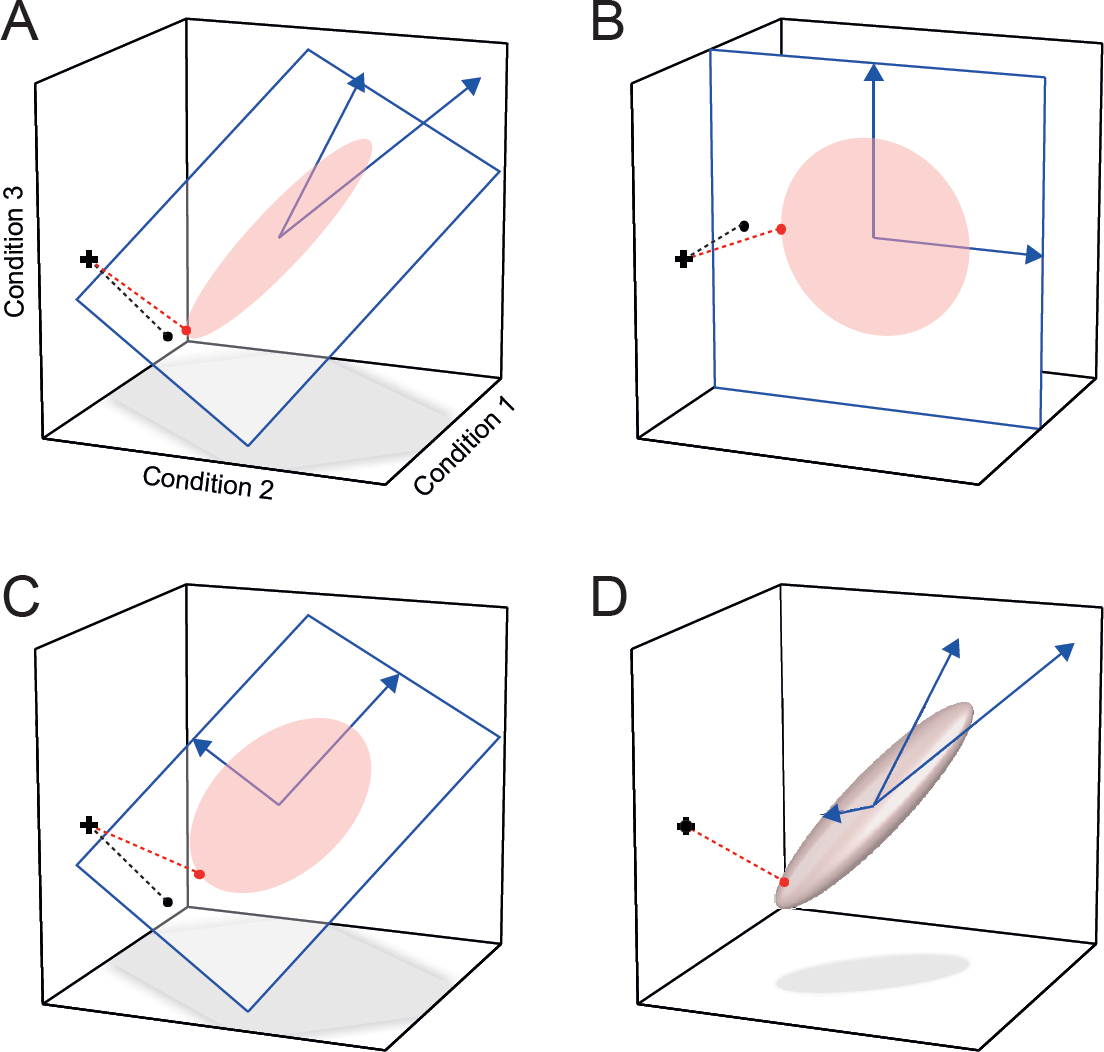
Adjudicating between encoding models with and without regularization. The axes of the three-dimensional space are formed by the response to three experimental conditions. The activity profile of each unit defines a point in this space. Models are defined by their features (blue arrows) and (when using regularization) a prior distribution of the weights for these features. The features and the prior, together, define a distribution of activity profiles (ellipsoids indicate an iso-probability-density contours of the Gaussian distributions). To predict the activity profile of a single measurement channel, the model is fitted to the training data set (cross). Simple regression finds the shortest projection (black dot) onto the subspace defined by the features, whereas regression with regularization (red dot) biases the prediction towards the predicted distribution. Two models (**A, B**) with features that span different model subspaces are distinguishable using regression without regularization. (**C**) This model spans the same subspace as model A. Unregularized regression results in the same projection as for model A, whereas regression with regularization leads to a different projection. (**D**) A saturated model with as many features as conditions. Unregularized regression can perfectly fit any data point (cross and black dot coincide). With regularization, the prediction is biased towards the predicted distribution (iso-probability-density ellipsoid).

Importantly, encoding analysis without regularization compares the subspaces of the competing models, but not their probability distributions. For example, the model depicted in Fig. 3C predicts a different distribution than the one in Fig. 3A. The features of these two models, however, span the same subspace. Therefore, without regularization, the predictions of these two models are identical (black dots) and the models indistinguishable.

#### Encoding analysis with regularization

When using regularized regression, encoding analysis evaluates models according to their predicted distribution of activity profiles. From a Bayesian perspective, regularization can be motivated by assuming a prior probability distribution on the weight vectors **w**_*.,i*_ the columns of **W**. Specifically, L2-norm (Tikhonov) regularization is equivalent to assuming a multivariate Gaussian prior with zero mean and variance-covariance matrix **Ω**. Under this assumption, the predicted second moment of the activity profiles is

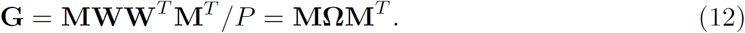

Thus, the model features together with the prior distributional assumption on the feature weights define a probability distribution over activity profiles. For example, a representational model of motor cortical activity could be defined by assuming that the features are individual units with cosine-tuning for different movement directions [18], and that (as a prior) the preferred directions of the units are uniformly distributed.

In practice, we allow a scalar factor, *s*, between the predicted and measured second moment. This accounts for the fact that different subjects or regions will have different signal levels and that hence the distribution of activity profiles have different widths. Under the assumption that the feature weights come from a multivariate Gaussian distribution with variance **Ω***s*, the best linear unbiased predictor (BLUP, [44]), i.e. the predictor that minimizes the squared error on the held-out data is:

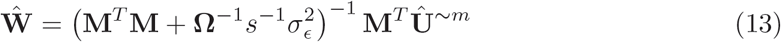
where 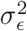 is the noise variance on the observations. The strength of regularization is determined by the ratio of this noise variance and the variance of the signal **Ω***s*, consistent with Bayesian inference of the weights on the basis of the prior and the data.

After assuming a prior on the model weights, the two models depicted in Fig. 3A and 3C predict different distributions of the activation profiles. When estimating the weights (Eq. 13), the activity profiles are projected onto the space spanned by **M**, but this time biased (red dot) towards the denser part of the model-predicted distribution of activity profiles. As a result, the two models make different predictions. An accurate prior will help the model generalize to the held-out data; an inaccurate prior will hurt generalization performance. The model with the distribution that is closest to the true distribution of activity profiles will yield the best cross-validation performance (as measured by *R*^2^ or *r*). When using regularized regression (Eq. 13), models can also have as many features as conditions (Fig. 3D), or even more features than conditions. When using unregularized regression, such *saturated* models are indistinguishable from each other. They become distinct only after adding weight-distributional priors.

Because regularization is equivalent to imposing a prior on the feature weights, it is not just a technical trick for estimation. Instead the prior is an integral part of the hypothesis being evaluated as it co-determines (together with the features) the probability distribution over activity profiles that the model predicts. Therefore, we will refer to encoding models evaluated using regularized regression analysis in the following as “encoding models with a prior”.

One important consequence of Eq. 12 is that the same representational model can be defined using different feature sets. Because a representational model is defined by its second moment, two feature sets **M**_1_ and **M**_2_, combined with corresponding second moment matrices of the weights, **Ω_1_** and **Ω_2_**, define the same representational model, if

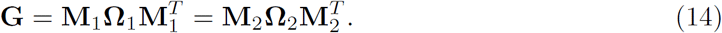

Thus, an important caveat when using encoding models is that one does not compare different feature sets per se – but rather different distributions (when using regularization) or different subspaces of activity profiles (when not using regularization). The winning model in either case can be equivalently re-expressed using a different feature set. Interpretation, therefore, must consider the model-predicted distributions or subspaces of activity profiles, not the particular feature basis set chosen (as the latter is not unique for any given representational model).

Technically, this also means that regression with a Gaussian prior can be implemented using ridge regression [45]. The equivalence is established by scaling and rotating the model matrix **M** in such a way that **Ω** becomes the identity matrix. Any representational model can be brought into this diagonal form by setting the columns of **M** to the eigenvectors of **G**, each one multiplied by the square root of the corresponding eigenvalue:

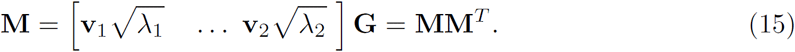

The strength of the regularization is determined by a scalar ridge coefficient defined by 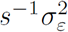. For an encoding model with regularization, the ridge coefficient still needs to be determined for each cross-validation fold. This can be done again by nested cross-validation [16], generalized cross-validation [46], or restricted maximum-likelihood estimation (Eq. 18). To save time, it is also possible to use a constant regularization coefficient. For our simulations, we estimated the optimal 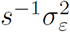 by maximizing Eq. 18 for the training set (across all voxels). Generalized cross-validation [46] yielded very similar results.

### Pattern component modeling

An alternative to cross-validation is to evaluate the likelihood of the measured activity profiles under the representational model. This approach is taken in pattern-component modeling [22]. We start with a generative model of the activity profiles (Eq. 6). We consider the activity profiles (columns of **U**) to come from a multivariate Gaussian distribution with zero mean and second-moment matrix **G**. To account for other nuisance effects (mean activity for each partition, low-frequency drift, etc), the model also has some fixed-effects regressors (**B**). We are not interested in fitting **U** per se, but simply want to evaluate the likelihood of the data under different models, marginalized over all possible values of **U**. The marginal distribution for each channel (columns of matrix **Y**) takes the form of a multivariate normal:

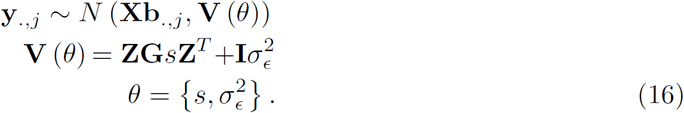

The predicted covariance matrix of the activity measurements for each person is the function of the model (as encoded in the second-moment matrix **G**) and two second-level parameters (*θ*): one that determines the strength of the signal (*s*) and one that determines the variance of the noise (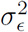). In determining the likelihood, we remove the fixed effects using the residual forming matrix

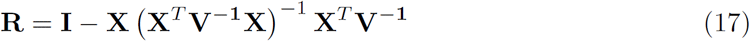

We need to then account for the removal of these fixed effects by evaluating the restricted likelihood *l*(**Y**|**G**, *θ*) [47]:

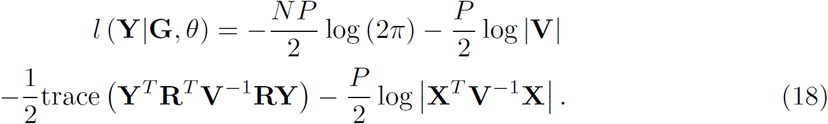

To evaluate the fit of a model, the scaling and noise parameters need to be determined. For fMRI data, these two parameters can vary widely between different brain regions and individuals, and are not meaningful in themselves. We therefore replace *θ* with point estimates that maximize Eq. 18 – i.e., the approach uses Empirical Bayes, or Type-II maximum likelihood for model comparison [45]. Because every model has the same two free second-level parameters, even models that are based on different numbers of features can be compared directly. An efficient implementation of this algorithm can be found in the open-source Matlab package for PCM [48].

### Representational similarity analysis

#### Relationship between representational dissimilarities and second-moment matrices

In RSA, representational models are conceptualized in terms of the dissimilarities between the activity patterns elicited across channels by the experimental conditions (Fig. 3C). One important dissimilarity measure is the Euclidean distance, which is closely related to the second-moment matrix **G**. The squared Euclidean distance between the true activity patterns for condition *i* and *k* (normalized by the number of measurement channels) is

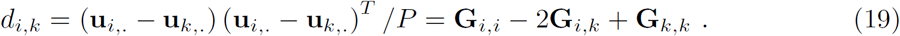

The Euclidean distance matrix is therefore a function the second moment of the activity profiles. The generalization of the Euclidean distances to non-isotropic noise is the Mahalanobis distance (see below). Correlation distances, another class of popular dissimilarity measures, can also be computed from the second-moment matrix. The cosine angle distance is defined as

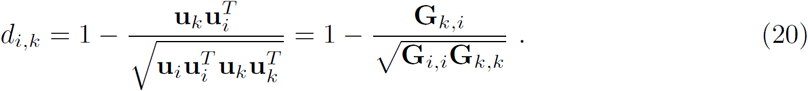

Here we focus on Euclidean and Mahalanobis distances, as they are independent of the resting baseline and generally easier to interpret [32].

In the following, we either represent these distances as a *K*×*K* representational dissimilarity matrix (RDM) **D**, or a *K(K-1)/2* vector **d** that contains all unique pairwise dissimilarities (the lower triangular entries of **D**). The vector of all pairwise dissimilarities can be obtained from **G** by defining a contrast matrix **C**, with each row encoding one of the pairwise contrasts, with a 1 and a − 1 for the contrasted conditions and zeros elsewhere:

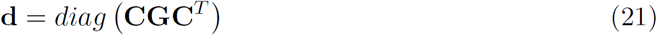

The distances contain the same information as the second moment matrix – however, we are losing the distance of each pattern to the baseline, which was encoded on the diagonal of **G**. Thus, in order to go from a distance matrix to a second-moment matrix, we need to re-set the origin of the coordinate system. An obvious choice is to define the mean activity pattern across all conditions to be the baseline. This is equivalent making the sum of all rows and columns of **G** zero, which can be achieved by defining the centering matrix **H** = **I**_*K*_ − **1**_*K*_/*K*, with **1**_K_ being a square matrix of ones. Under these conditions, **G** can be computed from **D** as

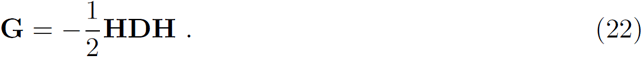

This yields the **G** that would be obtained if the patterns in **U** were centered about the origin, as can be achieved by subtracting the mean pattern from each pattern.

#### Multivariate noise normalization and cross-validation: the crossnobis estimator

A particularly useful dissimilarity measure is the cross-validated, squared Mahalanobis distance estimator (or crossnobis estimator for short). This estimator has superior characteristics in terms of reliability and interpretability as compared to other dissimilarity measures [32].

The crossnobis estimator uses multivariate noise normalization (see section Spatial dependence) to make the errors of different measurement channels approximately independent of each other. Euclidean distances (Eq. 19) computed on these pre-whitened activity estimates are equivalent to the Mahalanobis distance defined by the error-covariance matrix between channels (for details see [29, 32]).

The crossnobis estimator is cross-validated to yield an unbiased estimate of the Mahalanobis distance (assuming that the error covariance is correctly estimated). Conventional distances, which are non-negative by definition, are positively biased when estimated on noisy data: When one replaces the true activity patterns in Eq. 19 with their noisy estimates, the expected value of the Euclidean distance will be always higher than the true distances, because the noise terms are squared and summed. We can obtain an unbiased estimate of the true distance by computing the difference vectors between the two activity patterns from two independent data partitions and taking the inner product of the difference vectors. Thus, if we have *M* independent partitions, the crossnobis estimator can be computed using a leave-one-out cross-validation scheme:

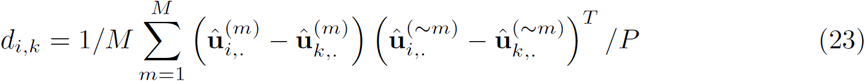
where 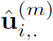 is the prewhitened pattern for condition *i* measured on partition *m*, and 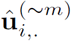 is same activity pattern determined from the data of all other partitions. The expected value of this estimator matches the true Mahalanobis distance [29, 32], except for a multiplicative bias caused by inaccuracies of the error covariance. In particular, if the patterns of two conditions only differ by noise, then the expected value of this measure will be zero. We will see below that the interpretable zero point can be advantageous for adjudicating among representational models.

#### Model comparison

In RSA, different representational models are evaluated by comparing the predicted to the observed dissimilarities. The overall magnitude of the Mahalanobis distances can vary considerably between subjects. The inter-subject variation is caused by differences in physiological responsiveness, physiological noise, and head movements – in short, by all the factors contributing to signal strength or the noise distribution, by which the Mahalanobis distance is scaled. Therefore, it is advisable to introduce a subject-specific scaling factor between observed and predicted distances, relying on the ratios between distances to distinguish models.

The unknown scaling of the observed dissimilarities is usually accounted for by calculating the correlation between the predicted and observed representational dissimilarity vectors (not to be confused with the use of correlation distance as an activity-pattern dissimilarity measure, Eq. 20).

The most cautious approach is to assume that we can only predict the rank ordering of distances [25]. It is then only appropriate to use Spearman correlation, or (in the case any of the models predict equal ranks for different pairs of conditions) Kendall’s *τ_a_* [27]. Evaluating models based on their ordinal dissimilarity predictions is conservative in terms of assumptions. However, the lesser reliance on assumptions comes at the cost of reduced sensitivity to certain differences between models. For more quantitative models, it may be appropriate to assume that distance predictions can be made on an interval scale. The assumption of a linear relationship between the predicted and measured distances motivates the use of Pearson correlation [28]. It may be justifiable in certain cases and can increase our sensitivity to differences between representational models.

Both rank-based and linear correlation coefficients not only allow an arbitrary scaling factor between observed and predicted distances, but also an arbitrary additive constant due the intercept of regression. However, the crossnobis estimator has an interpretable zero point: If a model predicts a zero distance for two conditions, then a brain region explained by the model should not be sensitive to the difference between the two conditions. This is a very meaningful prediction, which we can exploit to discriminate among models. Pearson and rank-based correlation coefficients discard this information. This suggest the use of a normalized inner product, a quantity analogous to a correlation coefficient, but in which the predictions and the data are not centered about their mean:

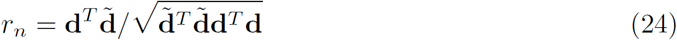

This amounts to a linear regression model between the predicted and observed distances, where the regression line is constrained to pass through the origin [49]:

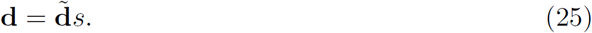

Here *s* is a scaling factor that is estimated from the data by minimizing the sum-of-squared errors between predicted and observed values.

Eq. 24 would provide optimal inference, if all distances estimates were independent and of equal variance. However, for the crossnobis estimator (and for most other dissimilarity measures), the assumptions of independence and equal variance are both violated. Estimated squared distances with larger true values are estimated with higher variability. Furthermore, the estimated distance between conditions A and B is not independent from the estimated distances between A and C [29]. To account for these factors, we need to know the predicted probability distribution of representational dissimilarity matrix estimates given a model. While the exact distribution of the vector of *K(K-1)/2* crossnobis estimates is difficult to obtain, we have shown that their distribution is well approximated by a multivariate normal distribution [29]:

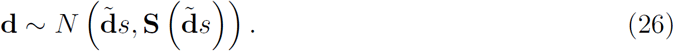

The mean of the distribution is the true distance matrix, scaled by a parameter relating to the signal strength in this subject (*s*). In [29], we showed that that the variance-covariance matrix of **d** is given by

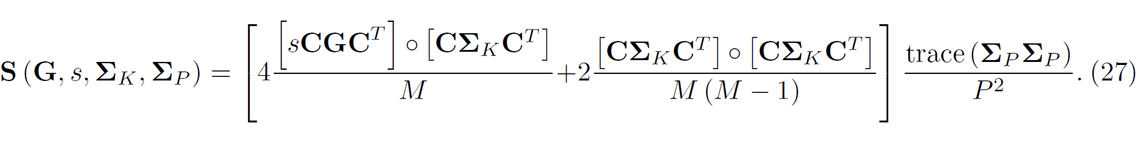

Where **G** is the predicted second-moment matrix of the patterns, **C** the contrast matrix that transforms the second-moment matrix into distances, and o refers to the element-by-element multiplication of two matrices. **Σ**_*K*_ is the condition-by-condition covariance matrix of the estimates of the activation profiles across partitions, which can be estimated from the variability of the activity patterns around their mean (Ū):

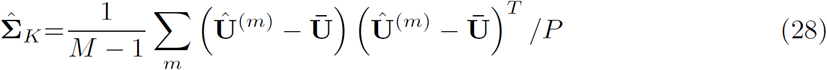

**Σ**_*P*_ is the voxel-by-voxel correlation matrix of the activation estimates. If multivariate noise-normalization [32] was completely successful, then this would be the identity matrix. However, given the shrinkage of the noise-covariance matrix used for noise-normalization, some residual correlations will remain; for accurate predictions of the variance, these must be estimated and accounted for [29].

Based on this approximation we can now express the log-likelihood of the measured distances **d** under the model predictions 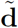.

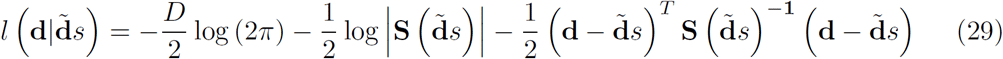

To evaluate the likelihood, we first need to estimate the scaling coefficient between predicted and observed distances by choosing *s* to maximize the likelihood. This can be done efficiently using iteratively-reweighted least squares (IRLS): Given a starting estimate of **S**, we can obtain the generalized least squares estimate of *s*,

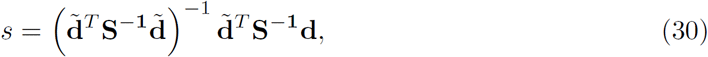
re-estimate **S** according to Eq. 27, and iterate until convergence.

### Simulated example data sets

We use simulated data sets here to evaluate and compare the three analysis techniques in a situation where the ground-truth is known. The three simulated example data sets are inspired by real fMRI studies. The first two examples are motivated by a paper investigating the representational structure of finger movements in primary motor and sensory cortex [28]. The structure of the empirically measured distances between movements of the five fingers was highly reliable across different individuals. The main question was whether this invariant structure is best explained by the correlations of finger movements in every-day life – i.e. the natural statistics of movement [50], or by the patterns of muscle activity required for these movements. Rather than hypothesizing that certain features form the basis set generating the activity profiles distribution, we could directly predict the second-moment matrices, and hence the RDMs, from the correlations between naturally occurring movements, or the correlations of muscle activity patterns. The predicted RDM for individuated movements of the five fingers (Exp. 1) is shown in Fig. 4A, B. The second example comes from experiment 3 in the same paper, this time looking at 31 different finger movements, which span the whole space of possible “piano-chord” combinations (Fig. 4C, D).

**Fig 4.**
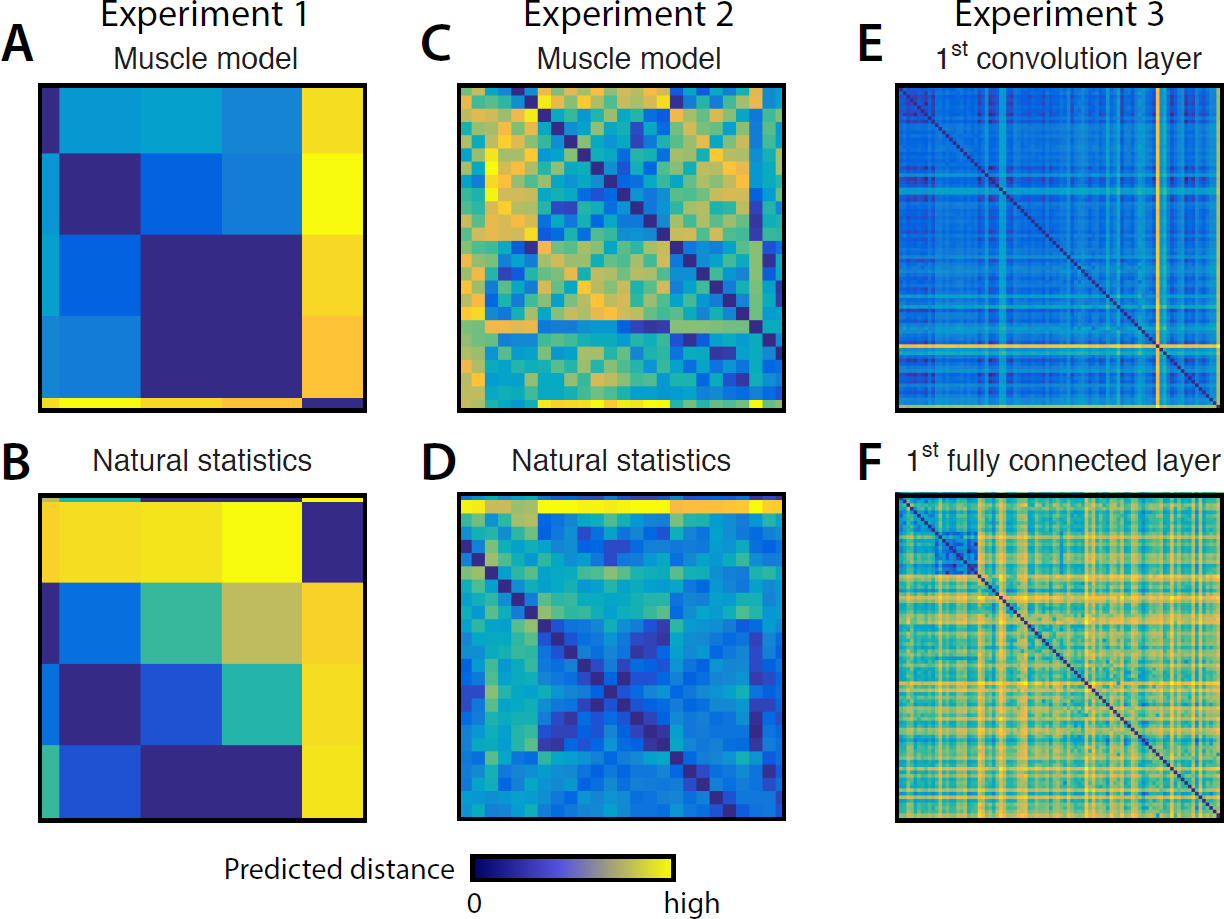
Representational dissimilarity matrices (RDMs) for the models used in simulation. Each entry of an RDM shows the dissimilarity between the patterns associated with two experimental conditions. RDMs are symmetric about a diagonal of zeros. Note that while zero is meaningfully defined (no difference between conditions), the scaling of the distances is arbitrary. For Experiment 1, the distance between the activity patterns for the five fingers are predicted from the structure of (**A**) muscle activity and (**B**) the natural statists of movement. In Experiment 2 (**C, D**) the same models predict the representational dissimilarities between finger movements for 31 piano-like chords. For Experiment 3 (**E, F**), model predictions come from the activity of the seven layers of a deep convolutional neural network in response to 96 visual stimuli. The 1st convolutional layer and the 1st fully connected layer are shown as examples.

The third example uses an experiment investigating the response of the human inferior temporal cortex to 96 images, including animate and inanimate objects [9]. The model predictions are derived from a convolutional deep neural network model – with each of the 7 layers providing a separate representational model. The bitmap images were presented to the deep neural network and the internal activity patterns used as representational models.

All data sets where simulated with 8 runs, 160 voxels, and independent noise on the observations. The noise variance was set to *σ*^2^ = 1. We first normalized the model predictions, such that the norm of the predicted squared Euclidean distances was 1. We then derived the second moment matrix (**G**) for each model using Eq. 22 and created true activity patterns that were normally distributed with second moment **UU**^*T*^/*P* = **G**_*s*_. The observation for each run were then generated by adding normally distributed random noise with unit variance to the data (Eq. 1). The signal-strength parameter *s* was varied systematically starting from 0 (pure noise data).

We generated 3,000 data sets for each experiment, parameter setting, and model. Each data set was generated by one model (ground truth) and was analyzed so as to infer the data-generating model, using each of the inference methods. To evaluate how well the methods adjudicated between the models, we compared the fit of the true model (i.e. the model that generated that particular data set) with each alternative model by counting the number of instances, in which the method decided in favor of the correct model. Thus, even though there were 7 alternative models in Experiment 3, chance performance for the pairwise comparisons was always 50%. The percentage of correct decisions over all possible model pairs and simulation was used as a measure of model-selection accuracy.

## Results

Our simulations illustrate the three main points of this paper: (1) Encoding approaches only provide a powerful test of representational models when using regularization that defines a prior distribution on the feature weights. (2) For the best possible inference using RSA, it is important to take the unequal variances and covariances between he distance estimates into account. (3) While PCM performs optimal model selection if the model assumptions are met, the other two approaches provide close approximations to the theoretical maximum. We will now discuss these results in turn.

### Encoding analysis without regularization

When evaluating encoding models without using regularization, one compares the subspaces spanned by the respective model features. To make different models distinguishable, one typically needs to reduce the dimensionality of the model matrix **M,** for example by using only the eigenvectors with the *n* highest eigenvalues of the predicted second-moment matrix. The decision to use a given number regressors is somewhat arbitrary: For example, Leo et al. [21] used 5 “synergies” (i.e. principal components of the kinematic data of 20 movements), as these explained 90% of the variance of the behavioral data.

Here we explore systematically how the number of principal components influences model selection. For each experiment, we simulated data sets with a fixed signal-to-noise ratio (Exp. 1 and Exp. 3: *s* = 0.3, Exp 2: *s* = 0.1; 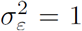), and compared model selection accuracies using a number of principal components ranging between one and the maximum number. We used both cross-validated performance measures, 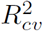 (Eq. 10) and *r* (the correlation between predicted and observed values; Eq. 11) to perform model selection.

Fig. 5A-C shows the percentage of correct model selections for Experiments 1-3. Results for encoding analysis without regularization are shown in blue. The dimensionality that differentiated best between competing models was 2, 3, and 5 features, respectively. As more features were included, the number of correct model selections declined. When the number of features was the same as the number of conditions minus 1 (due to the mean subtraction), i.e. the models became saturated, model selection accuracy fell to chance. This is expected, as two saturated models span exactly the same subspace and hence make identical predictions (Fig. 3D).

**Fig 5.**
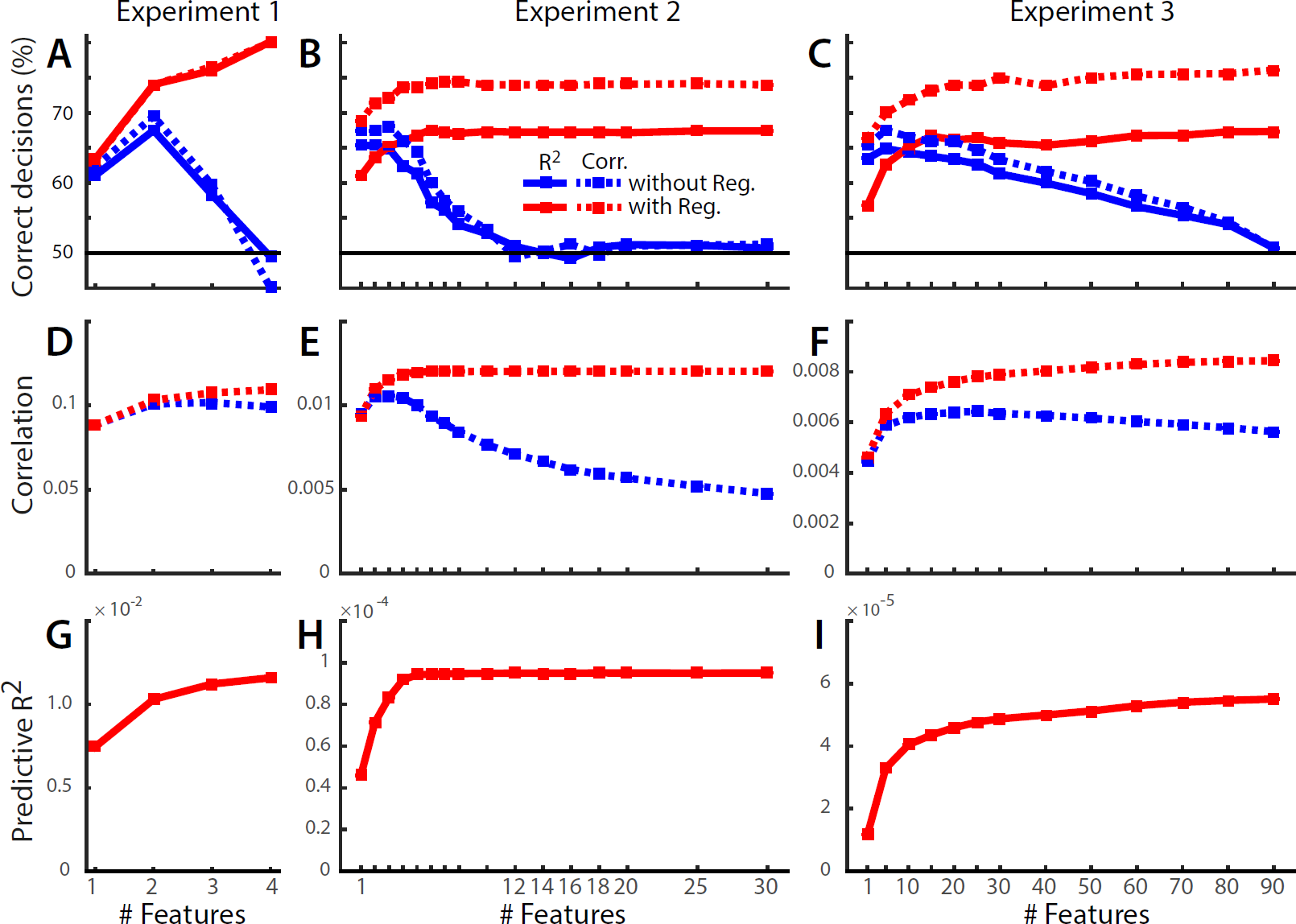
Dependence of encoding model analysis on regularization and the number of included model features. (**A-C**) Percent correct model selections using either 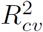 (solid line) or correlation (dashed line) for encoding models without a prior (blue lines) and with a prior (red line). (**D-F**) Correlation between predicted and observed patterns. (**G-I**) Predictive 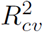 for the encoding models with prior. All 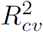 values for models without prior are negative, and therefore not visible.

Using correlations as selection criterion led to more accurate decisions than using 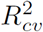. Correlations (Fig. 5D-F, blue lines) were generally positive and peaked at a number of features that was slightly higher than the optimal dimensionality for model selection. 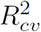 values for encoding without a prior were all negative (and are therefore not visible), because the approach does not account for the noise in the data and hence leads to predictions that are too extreme – i.e. the approach over-predicts the scale of the data. Correlations are insensitive to this problem as they allow for arbitrary scaling between predicted and observed values.

### Encoding approaches with regularization

From a Bayesian perspective, employing regularization (Eq. 13) is equivalent to adding a prior to the feature weights. Note that this changes the representational hypotheses tested. For example, the models for Experiment 3, based on the neural network representations, now predicted not only that some weighted combination of the neural network features can account for the data, but more specifically that the distribution of activity profiles should match the distribution of activity profiles of the original neural network simulation. In the model matrix, we scaled each principal component of **G** with the square root of the eigenvalue (Eq. 15), such that we could employ ridge regression to obtain the best linear unbiased predictor for the held-out data patterns.

For encoding models with a prior, model selection performance increased with increasing number of features (red lines, Fig. 5A-C). Thus, dimensionality reduction of the model is not necessary here. Furthermore, model selection was always more powerful with than without a prior when correlation was used for model selection. This reflects the fact that the prior provides additional information about the models to be compared. It enables us to compare well-defined distributions of activity profiles instead of just subspaces.

For Experiments 2 and 3, the 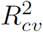 criterion performed substantially worse than the correlation between predicted and observed activity patterns. The difference between the two criteria arises from the fact that correlations allow for an arbitrary scaling between predicted and observed activity patterns, whereas 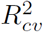 penalizes deviation in scale. The scaling of the prediction in turn strongly depends on the choice of the scalar regularization coefficient. This fact is illustrated in Fig. 6, where we simulated data from Exp. 2 with a fixed noise and signal strength, and varied the regularization coefficient systematically. While 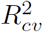 is highly sensitive to the choice of the regularization coefficient, the correlation criterion is not. Because the regularization coefficient is determined separately for each cross-validation fold and model, deviations from the optimal ridge will decrease model selection accuracy for the cross-validated 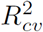 criterion, but not for the correlation criterion.

**Fig 6.**
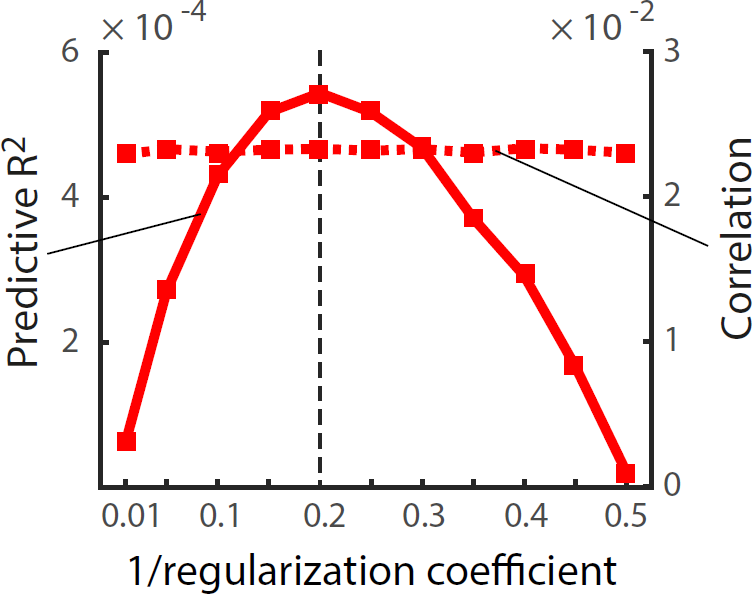
Sensitivity of the 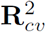 (solid line) and correlation (dashed line) to the choice of the regularization coefficient. Simulations come from Experiment 2 with a true signal strength of s=0.2 and a noise variance of 1. For this combination the optimal regularization coefficient is 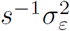 (dashed vertical line). The correlation criterion is generally robust against non-optimal choice of regularization coefficient.

In sum, using regularization improves model selection performance, even if the encoding model has fewer features than conditions or measurements. Rather than just comparing subspaces, the implicit prior on the weights means that a more specific hypothesis is being tested. From this perspective, it is unsurprising that we can adjudicate between these hypotheses with greater accuracy. Furthermore, the use of correlation instead of the predictive 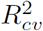 makes model selection more robust against variations in the regularization coefficient.

### Representational similarity analysis

When evaluating models with RSA, there is no need to restrict the model to a specific number of features – the second-moment matrix from all features can determine the predicted distances. As an empirical dissimilarity measure, we used the crossnobis estimator [32] and compared the predicted to the measured RDM. To select the winning model, we used rank-based correlation of dissimilarities [27], Pearson correlation, correlation with a fixed intercept (Eq. 24), and the likelihood of the observed distances under the normal approximation (Eq. 26) using the full variance-covariance matrix of the estimated dissimilarities.

For Experiment 1 (Fig. 7), rank-based correlation performed substantially worse than the other criteria. The lower performance of rank correlation may have been exacerbated here by the fact that the two models predict relatively similar dissimilarity ranks. However, we expect lower performance for rank correlation in general, because this approach does not use all the information in the measured RDMs. It forgoes the assumption of a linear relationship between predicted and measured dissimilarities and therefore does not exploit the information in the continuous magnitudes of the dissimilarities. Likelihood-based RSA yielded the best decisions; slightly better than Pearson correlation and fixed-intercept correlation.

**Fig 7.**
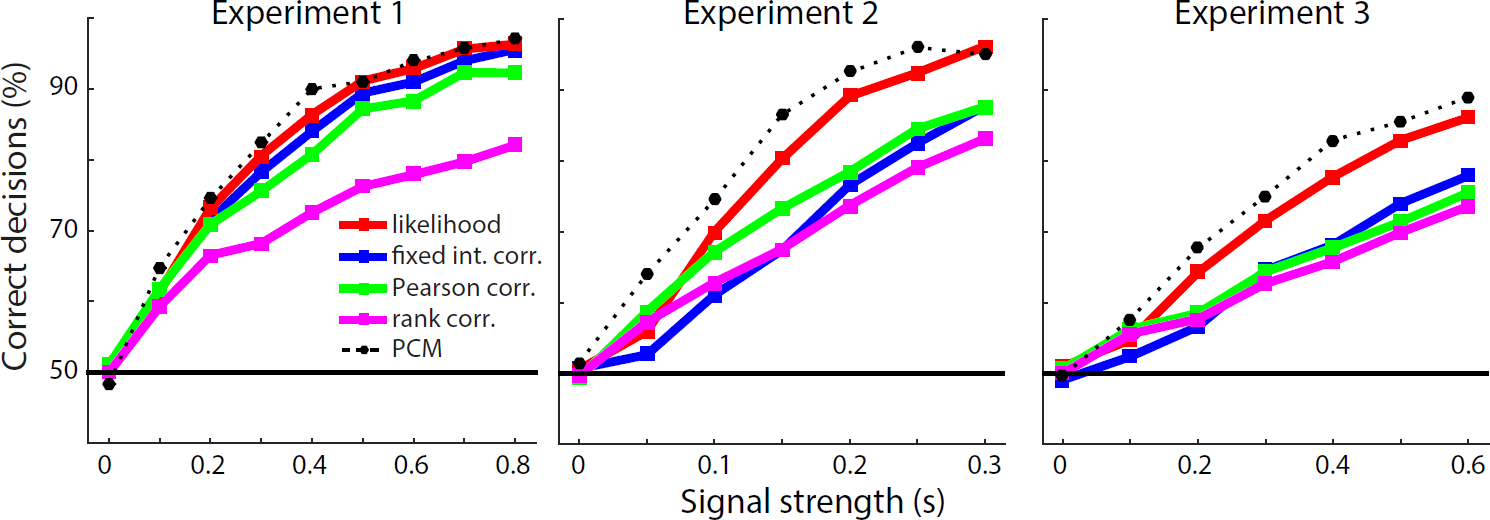
RSA model selection accuracies for different criteria of RDM fit. Data sets for all three experiments were generated with varying signal strength (horizontal axis). The percentage of correct decisions using different criteria is shown (dotted line). Models were selected based on the Spearman rank correlation (purple), Pearson correlation (green), fixed intercept correlation (blue) or likelihood under the multinormal approximation (red). For comparison, the model selection accuracy for PCM is shown in the dotted line.

The advantage of the likelihood-based approach was clearer for Exp. 2 and 3. Here, it led to about 10 percentage points greater accuracy of the decisions than the next-best RSA approach. This advantage is likely due to the fact that Pearson correlations and especially fixed-intercept correlations (Eq. 24) are sensitive to the observed value for the largest predicted dissimilarities, as these data points have a large leverage on the estimated regression line. Indeed, some of the models for Exp. 2 and 3 contain a few especially large dissimilarities, which will influence the model fit strongly. The likelihood-based approach incorporates the knowledge that large dissimilarities are measured with substantially larger variability [29], and hence discounts their influence. Notably, rank-based correlation performed relatively well on these models as compared to Pearson correlation, likely because rank correlation is robust to outliers and less dominated by the large predicted distances.

In sum, these simulations show that it is advantageous to take the covariance structure of the measured dissimilarities into account whenever the additional assumptions this requires are justified.

### Pattern component modeling

In the same simulations, we also applied the direct likelihood-ratio test, as implemented by PCM. As all the assumptions of the generative model behind PCM are met in the simulation, we would expect, by the Neyman-Pearson lemma [24], that this method should provide us with highest achievable model selection accuracy. Model selection performance (dotted line in Fig. 7) was indeed systematically higher than for the best RSA-based method. For direct comparison of the so far best methods – PCM, likelihood-based RSA, and encoding analysis with regularization (using correlations as a model selection criterion) – we simulated the three Experiments at a single signal strength (Fig. 8).

**Fig 8.**
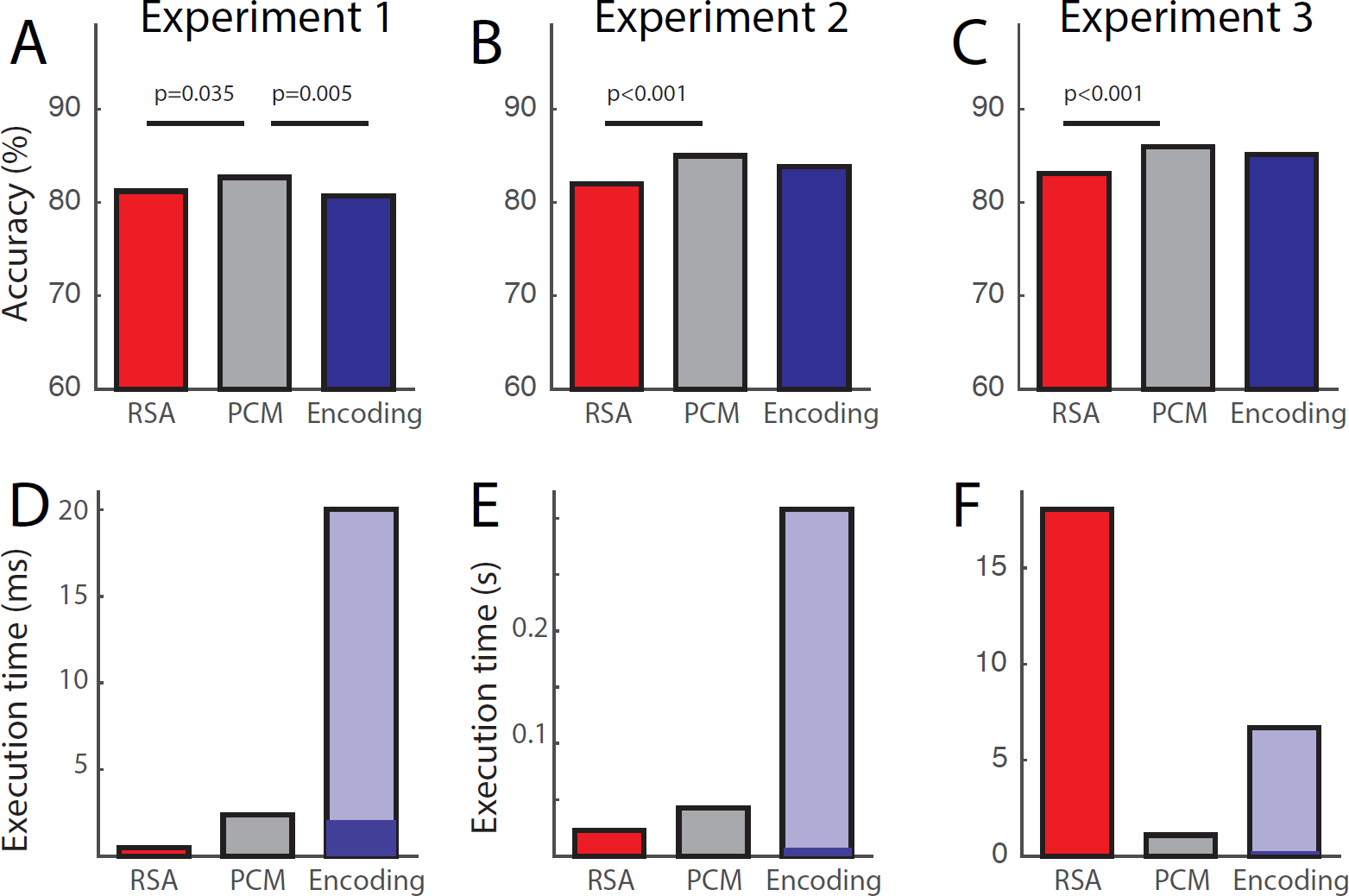
Model selection accuracy and execution time for likelihood-based RSA, PCM, and encoding analysis with regularization. (**A-C**) Model-selection accuracy was inferentially compared between the three techniques on the basis of N=3,000 simulations, using a likelihood-ratio test of counts of correct model decisions [51]. The signal-strength parameter for the simulation was set to s = 0.3 for Exp. 1, s = 0.15 for Exp. 2, and s = 0.5 for Exp. 3. All resulting significant differences (two-tailed, p<0.01, uncorrected) are indicated by a horizontal line above the bars. (**D-F**) Execution times for the evaluation of a single data set under a single model. For encoding, the time is split into the time required to estimate regression coefficients (dark blue) and the time to determine the regularization constant (light blue).

In this simulation, PCM resulted in 1.48, 3.01 and 2.86 percentage points (for Exp. 1-3, respectively) better model selection accuracy than likelihood-based RSA, and 1.98, 1.17 and 0.85 percentage points higher model selection accuracies than an encoding analysis using correlations. PCM never performed worse than another method and performed significantly better than the other two approaches in 4 of 6 total comparisons across the three experiments (Fig. 8). There were no significant performance differences between RSA and encoding analysis. Overall, however, all three methods were very close in performance.

### Computational cost

A practical concern is the speed at which the model comparison can be performed. This is usually not important when evaluating the model fit on a small number of participants or ROIs. However, if a larger number of models is evaluated continuously over the cortical surface using a searchlight approach [52, 53], or in data sets with large numbers of participants, computational cost becomes a practical issue. While we cannot treat this issue exhaustively, we provide here a brief overview over the computation time required for the three methods for our specific examples and implementation. In general, the computation time will of course depend on the number of conditions, the number of channels, the exact variant of each technique. Our goal here is simply to give the reader a starting point for making a choice for a particular application, trading off computational and statistical efficiency.

Both RSA and PCM operate on the inner product matrix of the activity estimates, thus the computational costs for these approaches is virtually independent of the number of voxels. PCM works on the *MK* × *MK* inner product matrix of the activity estimates, whereas RSA operates on a *K* × *K* matrix of distances between conditions. For a small number of conditions, this explains the favorable computational cost of RSA. However, when using likelihood-based RSA, the covariance matrix of the distances needs to be calculated and inverted. The size of this matrix is (*K*(*K*-1)/2)^2^ and it therefore grows with the 4^th^ power of the number of conditions *K*. For Exp. 3 (Fig. 8F) with *K* = 96, this is computationally costly, whereas PCM still only needs to invert matrices of size (*MK*)^2^. Using RDM-correlation-based model selection (whether rank, Pearson or fixed-intercept), RSA is much more computationally efficient (not shown).

For encoding models, conducting the actual ridge regression for each cross-validation fold (dark blue area) is extremely fast and efficient. The main cost of the technique lies in the determination of the optimal ridge coefficient (light blue area). In our simulations, we use restricted maximum likelihood estimation (Eq. 18) to do so – therefore this cost is always *M* times higher than for PCM alone. Depending on the implementation, generalized cross-validation [46] may offer a considerable speed-up. If very high speeds are required, one could use a constant ridge coefficient and accept the possible loss in model selection accuracy. In sum, while PCM is computationally feasible across the three experiments, encoding models were less efficient in the present implementation and likelihood-based RSA was less efficient than PCM for the condition-rich scenario of Experiment 3. Alternative variants of encoding models (with fixed ridge coefficient) and RSA (with correlation-based model selection) are less statistically efficient, but beat PCM in terms of computational efficiency.

## Discussion

In this paper, we defined representational models as formal hypotheses about the distribution of the activity profiles in the space defined by the experimental conditions. That is, a representational model specifies, which features are represented in a brain region, and how strongly they are represented. The “strength” of representation of a feature has two aspects: the number of responses (e.g. neurons) dedicated to a feature and the scaling of their activity profiles relative to the noise. The second-moment matrix of the activity profiles captures the combined effect of these aspects of feature strength. Two distinct representations with identical second-moment matrices therefore support linear decoding of any given feature at the same signal-to-noise ratio. This holds independent of the question whether the distribution of activity profiles is Gaussian. It motivates using the second moment as a summary statistic for characterizing representations. RSA, PCM and encoding models offer different tests of representational models, but all three depend, explicitly or implicitly, on the second-moment matrix to characterize each representational hypothesis. Thus, these methods are deeply related and should be understood as part of the same multivariate toolbox. The main characteristics of the three methods are summarized in Table 1.

### Encoding models without prior define subspaces, not distributions of activity profiles

There is a fundamental difference between encoding models with and without weight priors. Without a prior on the feature weights, encoding models test how well the subspace spanned by the model features captures the observed activity profiles. For models to be discriminable, the dimensionality (i.e. the number of features) of each model must be substantially lower than the number of experimental conditions. As the number of model dimensions increases, the subspaces of competing models increasingly overlap. Once the number of features matches the number of experimental conditions, their subspaces comprise the entire space of activity profiles, each perfectly fits the training data, and their predictions for unseen data become identical.

A subspace specifies what activity profiles are possible and what activity profiles are impossible (though they might still arise as estimates because of the noise). A subspace might be conceptualized as an infinite flat distribution over the subspace dimensions, with 0 probability outside the subspace. However, a uniform distribution on an infinite interval has an infinite second moment and hence does not specify the neural representation uniquely.

L2-norm regularization (i.e. ridge regression) is equivalent to imposing a Gaussian prior on the regression weights. With such a prior, the representational model specifies a probability distribution with a finite second moment. When changing the form of regularization, one also changes the implicit prior, and hence the representational model that is being tested. Thus, regularization is not simply a trick for stabilizing the fit. Instead, the weight prior forms an integral part of the model, which determines the strength with which each feature is encoded according to the model. Choosing a specific form of regularisation therefore constitutes a decision about the neuroscientific hypothesis to be tested rather than a methodological consideration.

### Encoding models tests hypotheses about activity profile distributions, not features sets

Encoding models do not support inferences about the particular feature set generating a representation, because infinitely many feature sets can span the same space. Even when using a prior, the feature set that characterizes a given representational model is not unique. Features should not in general be constrained to be orthogonal in the space of experimental conditions, because the structure of the model is not usually meant to depend on the experimental conditions chosen. Whether the features chosen are orthogonal or not, there is an infinite number of basis sets of features that express the same representational model (inducing the same second moment of activity profiles, Eq. 3). For example, two equally long correlated feature vectors can equally well describe a distribution with elliptical isoprobability-density contours (Fig. 3A) as two orthogonal features, with one vector longer than the other. Thus, when one representational model is shown to be superior to others, it does not imply anything special about the feature set chosen to express that model. These complications need to be kept in mind in the interpretation of the results of encoding model analyses. It is very tempting to attribute meaning to the particular feature basis chosen, especially when they are mapped onto the cortical surface [17, 21]. When interpreting these maps, one needs to remember that a feature set only describes a distribution of activity profiles, and that very different maps can emerge when the same distribution is described by a rotated set. In PCM and RSA, the equivalence of different feature sets is made explicit, as they lead to the same second-moment and representational dissimilarity matrices.

### Likelihood-based RSA is more sensitive than correlation-based RSA

When using RSA to test representational models, the crossnobis estimator provides a highly reliable measure of dissimilarity with the added advantage of having an interpretable zero-point [32]. Rank-based, Pearson, and fixed-intercept correlation provide fast and straightforward ways of measuring the correspondence between predicted and observed distances, so as to select the representational model most consistent with the data. However, using simple correlations ignores the dependence of the distance estimates, as well as their unequal variances. In other words, the sampling distribution of the estimated RDM in the space spanned by the dissimilarities (one dimension per pair of conditions) is not isotropic. This problem is addressed in likelihood-based RSA, which uses a multivariate-normal approximation to the sampling distribution of the crossnobis RDM estimate [29]. The approximation provides an analytical expression for the statistical dependency of distance estimates, as well as their signal-dependent variances. In the simulations, likelihood-based RSA was shown to be more powerful than correlation-based RSA. Its model-selection accuracy was only slightly below the theoretical upper bound, as established by PCM. Likelihood-based RSA might therefore become the approach of choice when comparing representational models using crossnobis estimates.

There are situations, however, in which the models are not specific enough to support ratio-scale predictions of representational dissimilarities. Moreover, for measurement modalities like fMRI, it might be undesirable to assume a linear relationship between predicted and measured representational dissimilarities. Rank-correlation-based RSA [25, 27] provides a robust method that is not dependent on the assumption of a linear reflection of the underlying neural dissimilarities in the data RDM. It is also more computationally efficient in the context of condition-rich designs. Likelihood-based RSA becomes computationally expensive as the number of conditions increases. A practical compromise might be to only use the diagonal of the variance-covariance matrix, which would dramatically reduce computational complexity at the expense of neglecting dependencies among dissimilarity estimates.

### Which method is best?

For all simulations, model selection using PCM [22] was better than competing methods. This is not surprising, as the data were simulated exactly according to the generative model underlying this approach (Gaussian distribution of noise and signal, independence across voxels). In this case, PCM implements the likelihood-ratio test, which by the Neyman-Pearson lemma [24] is the most powerful test. Beyond confirming what we know from theory, the simulations were important because they revealed how close the other two techniques come to the theoretical upper bound established by PCM. Results showed that encoding models with a prior and likelihood-based RSA perform near-optimally. In practice, we therefore expect these three techniques to provide similar answers. When its assumptions hold, PCM has clear advantages for model comparison, providing optimal power at reasonable computational cost. However, the other two techniques have other advantages that make them attractive for specific applications.

RSA using RDM correlation for model selection gives up statistical efficiency for computational efficiency, and beats PCM at the latter. When rank correlation is used to compare RDMs, the inference does not rely on a linear relationship between the true dissimilarities and the estimated dissimilarities, an assumption that might be violated in many contexts. RSA also provides readily interpretable intermediate statistics (cross-validated distances), which are closely related to linear decoders for all pairs of stimuli. These statistics can be used to test whether two conditions have different activity patterns [27, 29], or whether the dissimilarity is larger for one pair than for another pair of conditions. Multidimensional scaling of the stimuli on the basis of their representational dissimilarities also provides an intuitive visualization of the representational structure [25], which can be very helpful in the generation of novel representational hypotheses.

In contrast, PCM sometimes demands complicated approaches to answer simple questions: For example, to test the hypothesis that a difference between two conditions is encoded, one would need to fit one model that allows for separate patterns and one model that does not – and then compare the marginal likelihood of these models. Furthermore, PCM requires the noise to be explicitly modeled, whereas RSA removes the bias arising from noise through cross-validated distances.

Encoding analysis explicitly estimates the first-level parameters that describe the response for each individual voxel. This enables the mapping of the estimated features onto the cortical surface to study their spatial distribution [17, 21].

In sum, the three methods are deeply related in that they test hypotheses about the second moment of the activity profiles. However, each method constitutes a unique perspective on the data and supports different kinds of exploratory analyses. We view the methods as complementary tools that are part of a single coherent toolkit for analyzing representations.

### Single-voxel vs. multi-voxel inference

An important issue, which we have not touched upon so far, is whether to perform model comparison on single or multiple voxels. While RSA and PCM are usually applied to groups of voxels (such as for ROIs or searchlights), encoding models are often compared on the single-voxel level. This tendency, however, is not strictly inherent in methodological constraints: The searchlight approach for RSA and PCM can be reduced to single voxels, and encoding models can be combined with multi-voxel searchlights. Analyses with coarser granularity give up some spatial precision of the map in exchange for greater statistical power. Searchlight mapping boosts power (1) by locally combining the evidence, (2) by enabling the use of a multivariate noise normalization, and (3) by reducing the effective number of multiple comparisons [54]. There is no reason to assume that a single-voxel searchlight is always the optimal choice when balancing spatial precision and power. Based on our previous results [32], we expect that ignoring voxel dependencies will entail a loss of sensitivity when making inferences on representational models for regions of interest comprising multiple responses.

### Testing models without overfitting to the noise and to the sample of experimental conditions

Whenever a model is fitted using experimental data, its parameters will necessarily be overfitted to the data to some extent. Assessing the performance of a fitted model therefore requires independent test data. An important question is whether the test data should consist in independent measurements for the same experimental conditions or in measurements for a fresh sample of experimental conditions (e.g. a different sample of visual images). The simple answer is that it depends on the inference we would like to make. If our hypothesis is restricted to the present set of conditions (e.g. five finger movements), we need only account for overfitting to the noise in the data and require different measurements for the same conditions. If our hypothesis is about a population of conditions (e.g. all natural images), we need to account for overfitting to the condition sample and require measurements for an independent random sample of conditions from the population of conditions covered by our hypothesis.

However, overfitting only needs to be accounted for when the model being tested had parameters fitted in the first place. Encoding models always require independent test sets to account for the over-fitting of the first-level parameters of the representational model (feature weights). RSA and PCM, by contrast, rely explicitly on summary statistics of the responses. Therefore, only second-level parameters related to the strength of the signal and noise need to be fitted (see Table 1). Because the representational models considered here had the same number of such second- level parameters, they could be compared directly.

### What about decoding approaches?

Decoding is widely used in multivariate analysis of brain imaging data [11–13]. Can it serve us also as a tool for comparing representational models? While one can use standard decoding approaches to determine whether specific features are represented in an area or not, it does not lend itself to the comparison of full representational models (as defined here). Representational models determine (via the second moment matrix) the decodability of any linear feature, not just a restricted set of features. This is most obvious in RSA, where the RDM assembles all pairwise condition discriminabilities. It is of course possible to use decoding in the context of the methods considered here. For example, some studies have used encoding models to decode stimuli [15, 16, 21]. Decoding accuracy can then serve, instead of correlation or 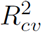, to evaluate the performance of an encoding model on held-out data. While this approach is motivated by the intuitive demonstration of mind reading, it does not provide a particularly natural or powerful approach to adjudicating between representational models. Alternatively, we could use classification accuracy as a measure of dissimilarity between two conditions in the context of RSA [55]. However, classification essentially converts a continuous measure of dissimilarity into a binary label of correct / incorrect. It is therefore expected to be less informative than the underlying continuous measure, and we have shown previously that this entails a loss of sensitivity in practice [32]. In sum, decoding is not particularly useful for the evaluation of representational models [14, 23] and should therefore be limited to situations, in which the quality of the decoding itself is the measure of interest.

### Flexible representational models

All models considered here were “fixed”, i.e., they did not include free parameters that would change the predicted second-moment matrix. In many applications, however, the relative importance of different features (for example encoding strength for orientation and color) are unknown. In this case, the predicted second moment can be expressed as the weighted sum of different pattern components, i.e. **G** = Σ_*i*_ *ω_i_***C_i_** [22, 56–58], with the weights being free second-level parameters. In other situations, **G** is a nonlinear function of free model parameters: For example, **G** depends non-linearly on the spatial tuning width in population receptive field modeling [59]. Both RSA and PCM already provide a mechanism to estimate such parameters, as both approaches already need to estimate the signal strength parameters *s* by maximizing the respective likelihood function (Eq. 17, 28) – and the analytical derivatives of the likelihood (Eq. 17, 28) with respect to the parameters are easily obtained. In the context of encoding approaches using ridge regression, free model parameters that change the model structure would result in independent scaling of different features, rotations, or extensions of the model matrix **M**. At the time of writing there are no published examples of such parameter optimization in the context of cross-validated encoding models that we know of.

The inclusion of free parameters into the model also enables the specification of measurement models. Representational models ideally test hypotheses about the distribution of activation profiles of the core computational elements – i.e. neurons. When using indirect measures of brain activity such as fMRI or MEG, the distribution of activity profiles across measurement channels is also influenced by the measurement process, which samples and mixes neuronal activity signals in the measurement channels [30, 60–63]. As the underlying brain computational models become more specific and detailed, the corresponding measurement models will also have to be improved.

### Higher-order moments of the activity profiles

We focused on approaches that characterize the distribution of activity profiles by its second moment. If the true distribution of the activity profiles is a multivariate Gaussian, then the second moment fully defines the distribution of activity profiles. However, a representational hypothesis may not only predict that the response for condition A is uncorrelated to the response for condition B, but, for example, that channels either respond to A or B, but not to both A and B. Such tuning is for example prevalent in primary visual cortex, where neurons (and voxels) respond to a stimulus in a *one* specific part of the visual field, but less often two or more disparate locations [59]. This would correspond to a non-Gaussian prior on the feature weights. In a recent publication, Norman-Haignere and colleagues [64] suggested a likelihood-based method, in which the Gaussian prior on the feature weights **W** is replaced with a Gamma distribution, essentially providing a non-Gaussian extension of PCM. It will be interesting to determine to what degree such non-Gaussian distributions are present in fMRI or single-cell data, and what computational function these may serve.

It is important to stress that the approaches considered here are still appropriate when the distribution of activity profiles is truly non-Gaussian. Even in the non-Gaussian case, the second moment determines the representational geometry and thus the decodability of all possible features. It therefore remains essential for characterizing the representation. Taking into account higher moments of the activity profile distribution would enable us to distinguish between representations that afford the same decoding of features (assuming that readout neurons have access to the entire code), but achieve this by distinct population codes.

### Conclusions

If advances in brain-activity measurements are to yield theoretical insights into brain computation, they need to be complemented by analytical methods to test computational models of information processing [65]. The main purpose of this paper was to provide a clear definition of one important class of models – representational models – and to compare three important approaches of testing these. We have shown that PCM, RSA and encoding analysis are all closely related, testing hypotheses about the distribution of activity profiles. Moreover, all three approaches, in their dominant implementations, are sensitive only to distinctions between representations that are reflected in the second moment of the activity profiles. Thus, these three methods are properly understood as components of a single analytical framework. Each of the three methods has particular advantages and disadvantages and preferred areas of application.

1. PCM provides an analytic expression for the marginal likelihood of the data under the model, and therefore constitutes the most powerful test for adjudicating between representational models if the assumptions hold. Its analytical tractability and relative computational efficiency are further attractive features, especially when considering models with increasing numbers of free parameters.
2. RSA provides highly interpretable intermediate statistics and is therefore ideally suited for the visualization and exploratory analysis. Furthermore, simple models are often more easily tested than with PCM. The normal approximation to the distribution of estimated distances enables inference that is nearly as powerful as the likelihood-ratio test provided by PCM. Finally, dissimilarity-rank-based RSA, though less sensitive, provides a means of inference that does not rely on the assumption of a linear relationship between predicted and measured dissimilarities and is computationally efficient even for condition-rich designs.
3. Encoding approaches enable the voxel-wise mapping of model features onto the cortical surface. They therefore are the natural choice when the spatial distribution of features or the voxel-wise comparison of representational models is the main interest.

We hope that the general framework presented here will enable researchers to combine these approaches to make progress revealing the computational mechanisms of biological brains.

